## Supplementary materials for "How individual vigor shapes human-human physical interaction"

#### 1 Solo session

##### 1.1 Effects of viscous resistance

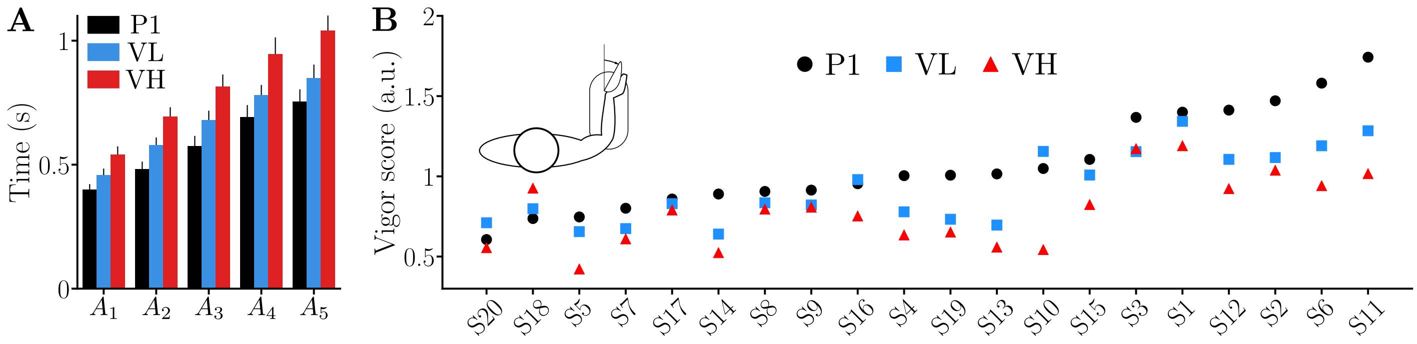


**Figure S.1: Effect of the viscous resistance on movement time and vigor. A.** Movement time increases with VL and VH compared to P1 and with VH compared to VL. **B.** The vigor scores of individuals in P1 are spread as in previous studies [1, 2]. VL induces a global decrease in vigor scores compared to P1 and VH induces a second reduction compared to VL.

##### 1.2 Minimum time-effort predictions in solo session

We simulated the behavior of the average participant in the solo session using a minimum time-effort (MTE) compromise (see Eq. 6 in the main text). This compromise was derived by identifying the average participant's cost of time through an inverse optimal control method applied to data from P1 [2–4]. The resulting model predictions for movement duration (MD) and the absolute work of the robot's viscous torque, in both VL and VH and for the five amplitudes, are summarized in Fig. S.2.

In shown in Fig. S.2, the MTE model accurately predicts both the duration of and mechanical work, yielding minimal prediction errors. These accurate predictions for the solo session make this model a valid basis for the model of the dyad session, where interaction with the partner can be interpreted as a modification of the cost of effort.

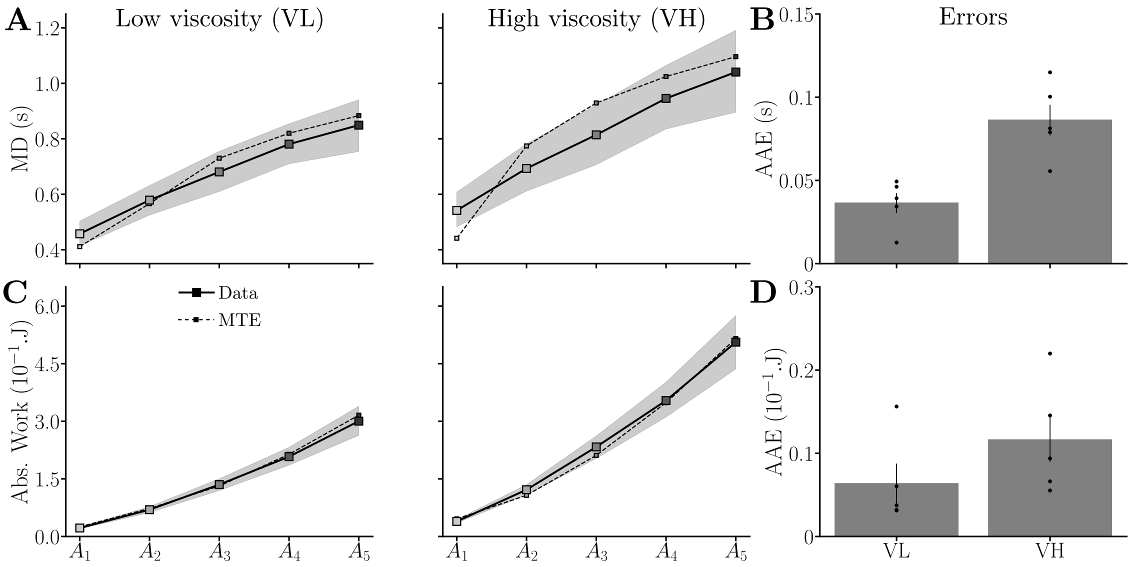


**Figure S.2: Minimum time-effort (MTE) compromise in the VL and VH conditions.** For both conditions, the behavior of the average participants is compared to an optimal time-effort compromise strategy (dashed line with markers). **A.** Predicted movement duration in the VL and VH conditions. **B.** Average absolute error (AAE) in terms of movement duration. **C.** Predicted absolute work of the robot torque in the VL and VH conditions. **D.** Average absolute error (AAE) in terms of absolute work.

### 2 Dyad session

#### 2.1 Complementary results on dyadic behavior

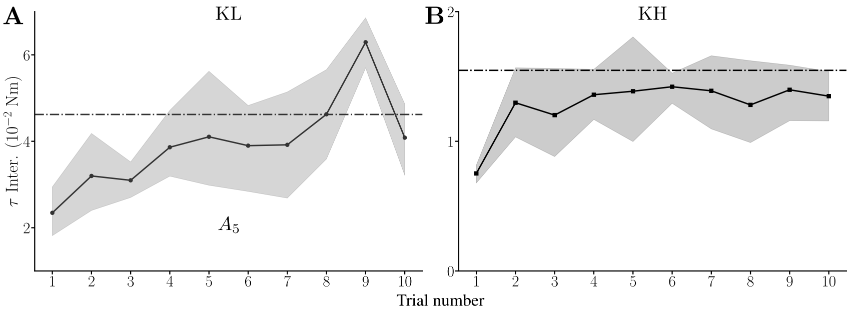


**Figure S.3: Adaptation of interaction torque through trials in the connected session.** The trends are computed using the first connected block performed by the dyads. **A.** First 10 trials of the largest amplitude in KL condition. The dashed line corresponds to the average behavior for the remaining trials towards this target. **B.** First 10 trials KH condition, as allowed by the absence of differences between targets for this block. The dashed line correspond to the average behavior during the rest of the KH block.

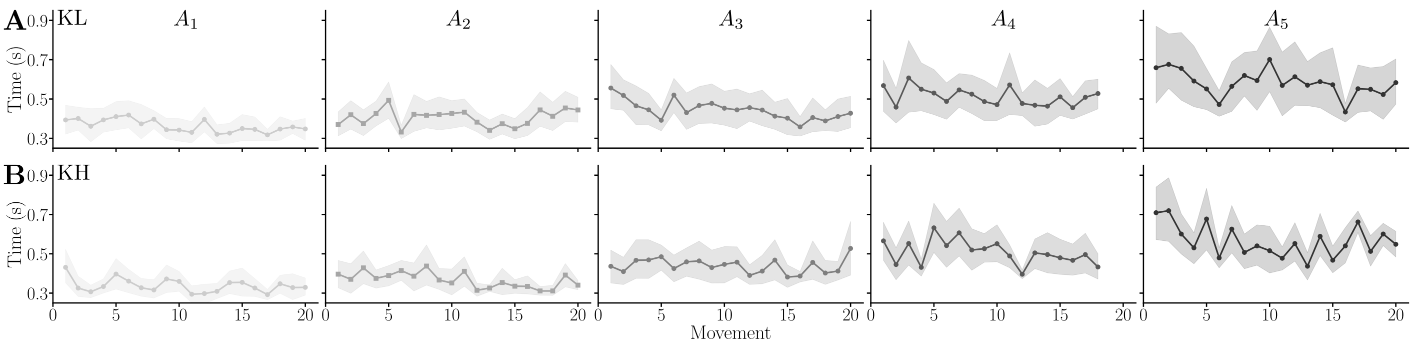


**Figure S.4: Limited adaptation of movement duration through trials in the connected session.** The trends are computed using the first connected block performed by the dyads. Columns correspond to the five targets. **A.** KL condition. **B.** KH condition.

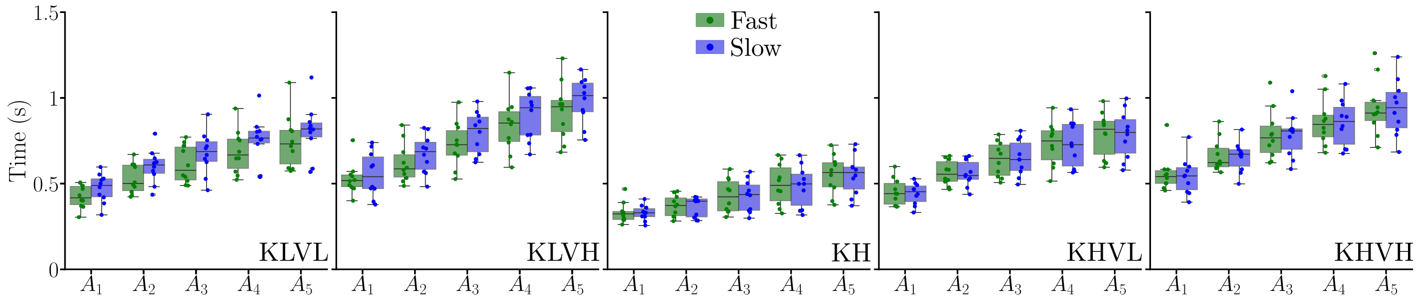


**Figure S.5: The fast and slow participants have similar movement durations when connected.**

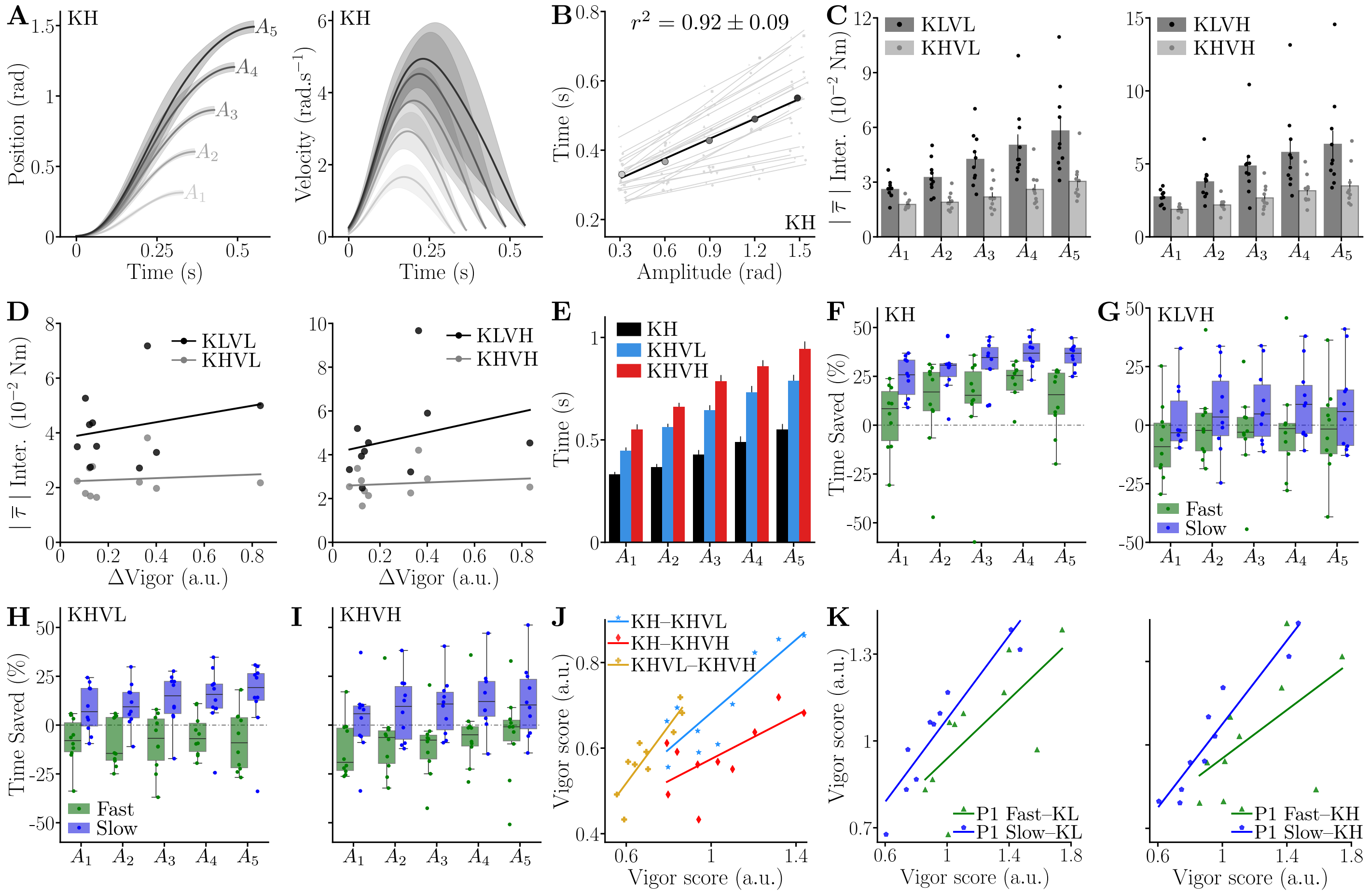


**Figure S.6: Complementary behavioral data of the dyadic session. A.** Trajectories and velocity profiles averaged across the population for KH and for the five targets. **B.** Individual and averaged amplitude-duration relationships during KH. **C.** Absolute average interaction efforts in presence of a viscous load. **D.** Correlation of the difference in solo vigor between partners and interaction efforts with a viscous load. **E.** Effects of the viscous load movement duration with KH. **F.** Percentage of time saved by the fast and slow groups between P1 and KH, positive values indicate faster movements during KH. **G.** Percentage of time saved by the fast and slow groups between VH and KLVH. **H.** Percentage of time saved by the fast and slow groups between VL and KHVL. **I.** Percentage of time saved by the fast and slow groups between VH and KHVH. **J.** Significant correlations of vigor scores between the three KH conditions. **K.** Examples of correlations between the vigor scores obtained by the fast and slow group during P1 and those obtained during KL and KH. Dyadic vigor is predicted by the vigor of the slow group (LMM, see main text).

#### 2.2 Complementary results on performance

**Performance** As we have shown when analyzing the movement duration of dyads, there was an increase in time performance for KL and KH, which was not observed when an additional viscous effort was imposed by the exoskeletons. However, it is yet unclear whether this reduction of movement time comes at the cost of a reduced accuracy as would predict Fitts' law [5], and whether it would impact other objective metrics of human movement performance such as smoothness [6]. Therefore, we conducted an analysis of movement smoothness and an analysis of accuracy. Accuracy was estimated by computing the standard deviation of the wrist joint position after the end of the main movement (see Methods for the segmentation of movements). The results of these analyses are summarized in Fig. S.7 alongside the saved time when compared to P1.

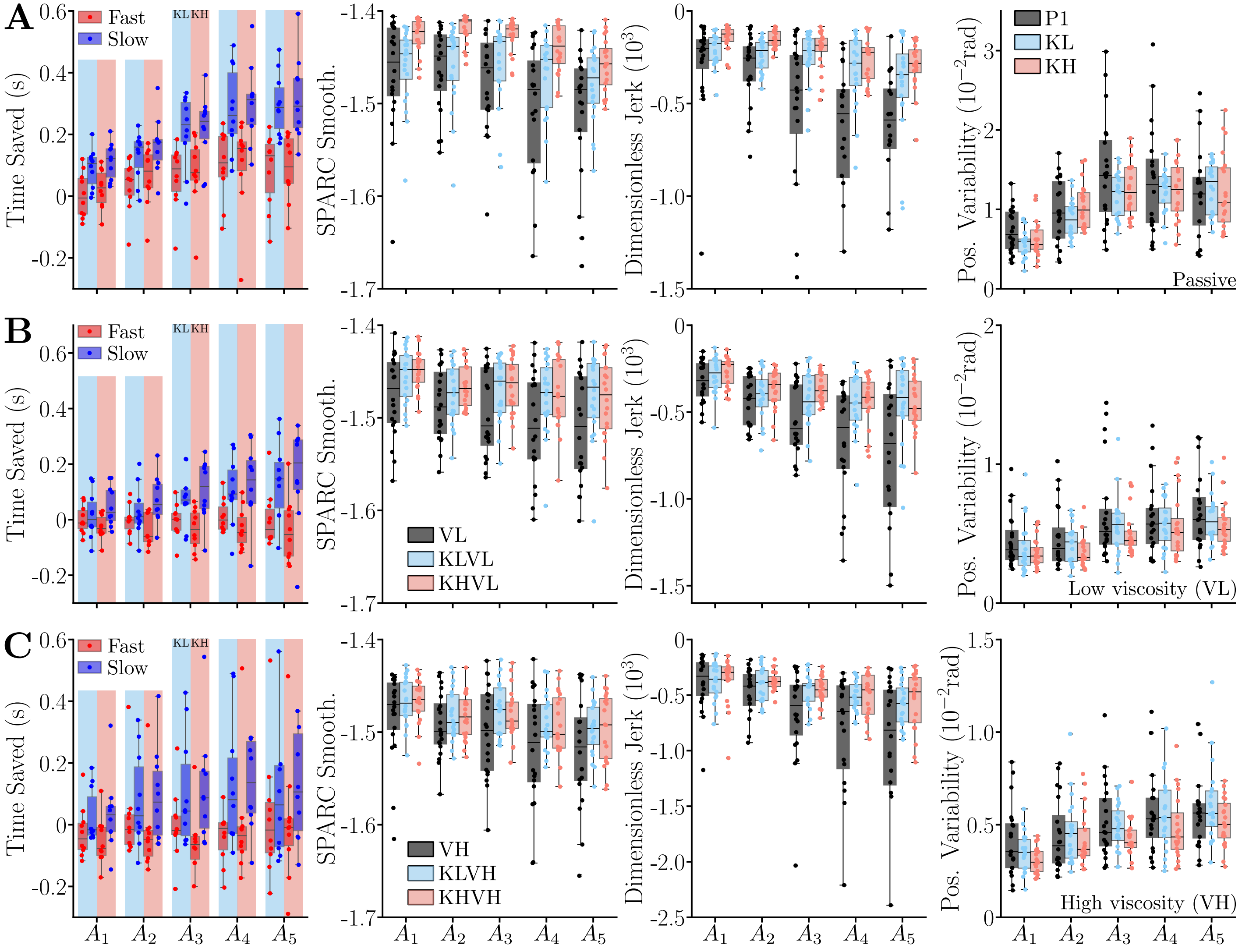


**Figure S.7: Movement performance of the dyads.** Metrics include the time saved when compared to the solo session, the SPARC smoothness of velocity profiles, the dimensionless jerk, and the standard deviation of position during stabilization (after the movement). All performance metrics are computed separately for the low (KL) and high (KH) connection stiffness. The time saved when compared to the solo session is split between the faster and slower participant. **A.** Passive conditions. **B** Low viscosity (VL) conditions. **C.** High viscosity (VH) conditions.

*Movement smoothness:* As highlighted by Fig. S.7 there is a trend of improved movement smoothness with both used metrics and for all conditions. Interestingly, a main effect of the human-human connection on movement smoothness was observed for all conditions, whether with passive exoskeletons (for both metrics: $W > .52$, $p < 10^{-3}$), with VL (for both metrics: $W > 0.23$, $p < 0.01$), or with VH (for the dimensionless jerk: $W = 0.23$, $p = 0.01$).

For the passive exoskeletons conditions (P1, KL and KH), the connection induced significantly smoother movements in KL and KH than in P1 according to both metrics (in all cases: $p < 2{\cdot}10^{-3}$, Cohen's $D > 0.83$), except SPARC for the comparison between P1 and KL. Furthermore, movements were smoother in KH than in KL (for both metrics: $p < 210^{-3}$, Cohen's $D > 0.58$).

For the low viscous resistance conditions (VL, KLVL and KHVL), the connection induced significantly smoother movements in KLVL and KHVL than in VL according to both metrics (in all cases: $p < 6{\cdot}10^{-3}$, Cohen's $D > 0.61$). No difference was observed between KLVL and KHVL.

For the high viscous resistance conditions (VH, KLVH and KHVH), the connection induced significantly smoother movements in KLVH and KHVH than in VH according to the dimensionless jerk (in all cases: $p < 0.025$, Cohen's $D > 0.49$). No difference was observed between KLVH and KHVH.

In sum, our analysis showed a clear improvement of movement smoothness for both KL and KH compared to when participants are moving alone. This result was shown to hold throughout five movement amplitudes and for three different task dynamics.

*Accuracy:* As highlighted by Fig. S.7 the reported reduction of movement time and increased smoothness did not seem to come at the expense of accuracy. In fact, the connection between participants had no significant effect on movement accuracy.

In sum, overall movement quality – in terms of time, smoothness, and accuracy – was either improved or not impacted (for the accuracy mainly) by the connection between participants. Importantly, this result was shown to hold for two levels of stiffness, throughout five amplitudes, and throughout three task dynamics. It can also be noted that movement accuracy was improved by the introduction of a viscous resistance in the dynamics of the exoskeletons, although the results, out of our scope of analysis, are not detailed here.

#### 2.3 Complementary results on interaction efforts

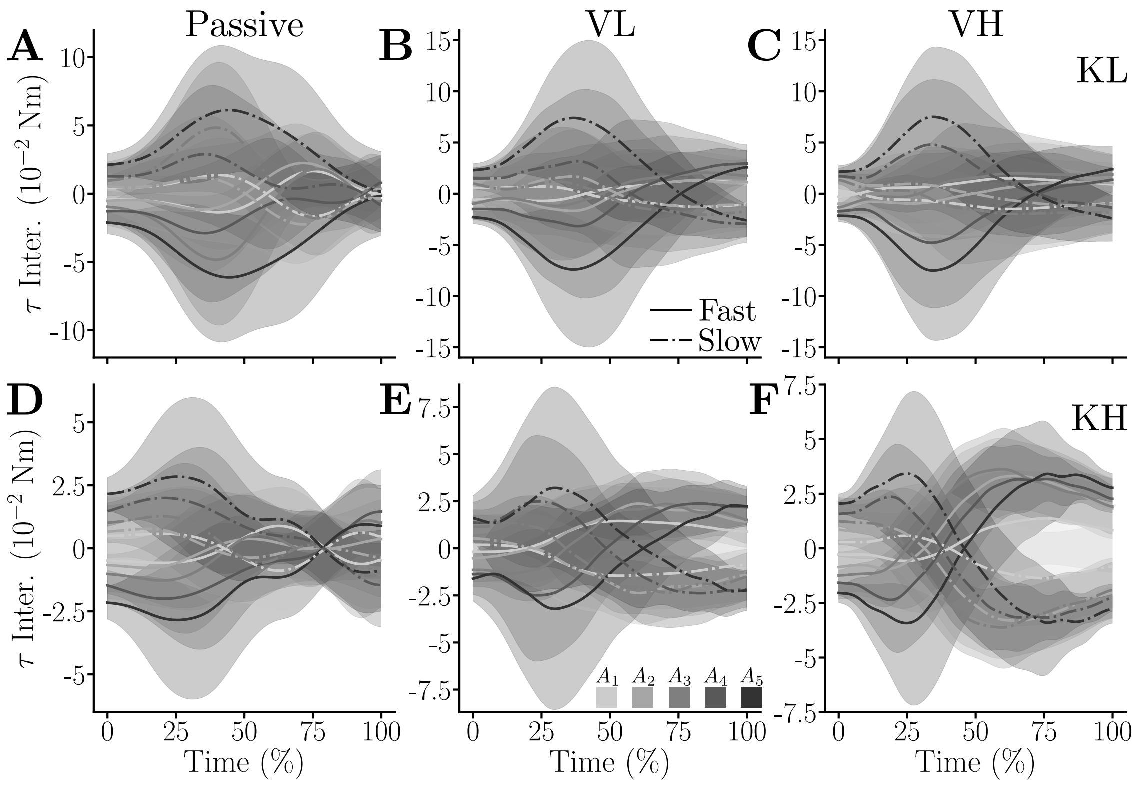


**Figure S.8: Average interaction efforts throughout conditions and targets for dyad D1.** Efforts received by the the more vigorous participant are represented with solid lines and those received by the less vigorous participant are represented with dashed-dotted lines. **A.** Low stiffness and passive robot (KL). **B.** Low stiffness and low viscous torque applied by the robot (KLVL). **C.** Low stiffness and high viscous torque applied by the robot (KLVH). **D.** High stiffness and passive robot (KH). **E.** High stiffness and low viscous torque applied by the robot (KHVL). **F.** High stiffness and high viscous torque applied by the robot (KHVH).

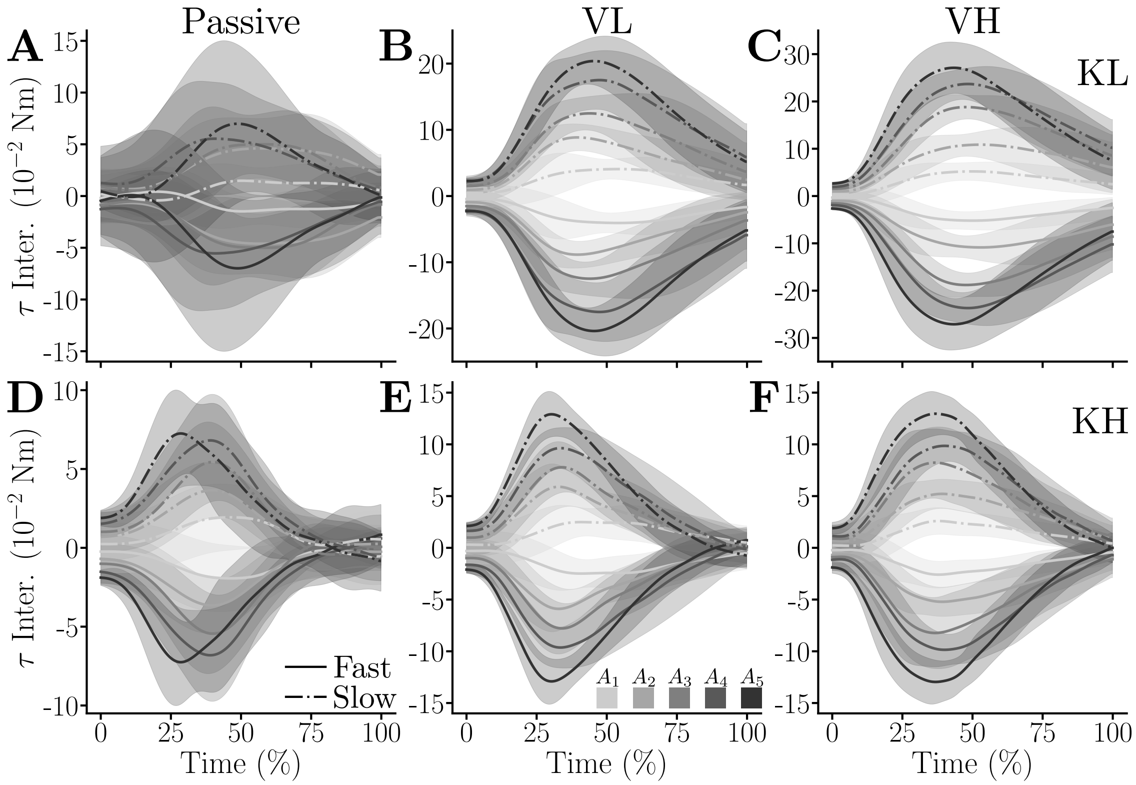


**Figure S.9: Average interaction efforts throughout conditions and targets for dyad D2.** Efforts received by the the more vigorous participant are represented with solid lines and those received by the less vigorous participant are represented with dashed-dotted lines. **A.** KL. **B.** KLVL. **C.** KLVH. **D.** KH. **E.** KHVL. **F.** KHVH.

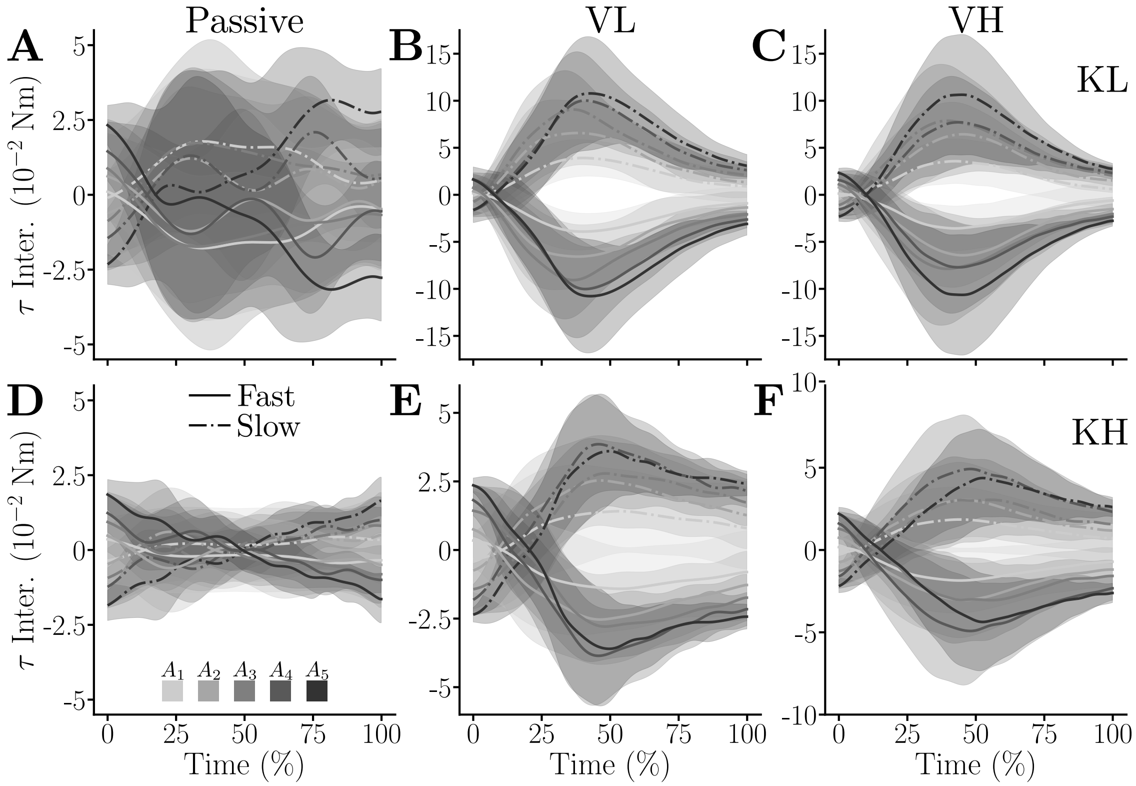


**Figure S.10: Average interaction efforts throughout conditions and targets for dyad D3.** Efforts received by the the more vigorous participant are represented with solid lines and those received by the less vigorous participant are represented with dashed-dotted lines. **A.** KL. **B.** KLVL. **C.** KLVH. **D.** KH. **E.** KHVL. **F.** KHVH.

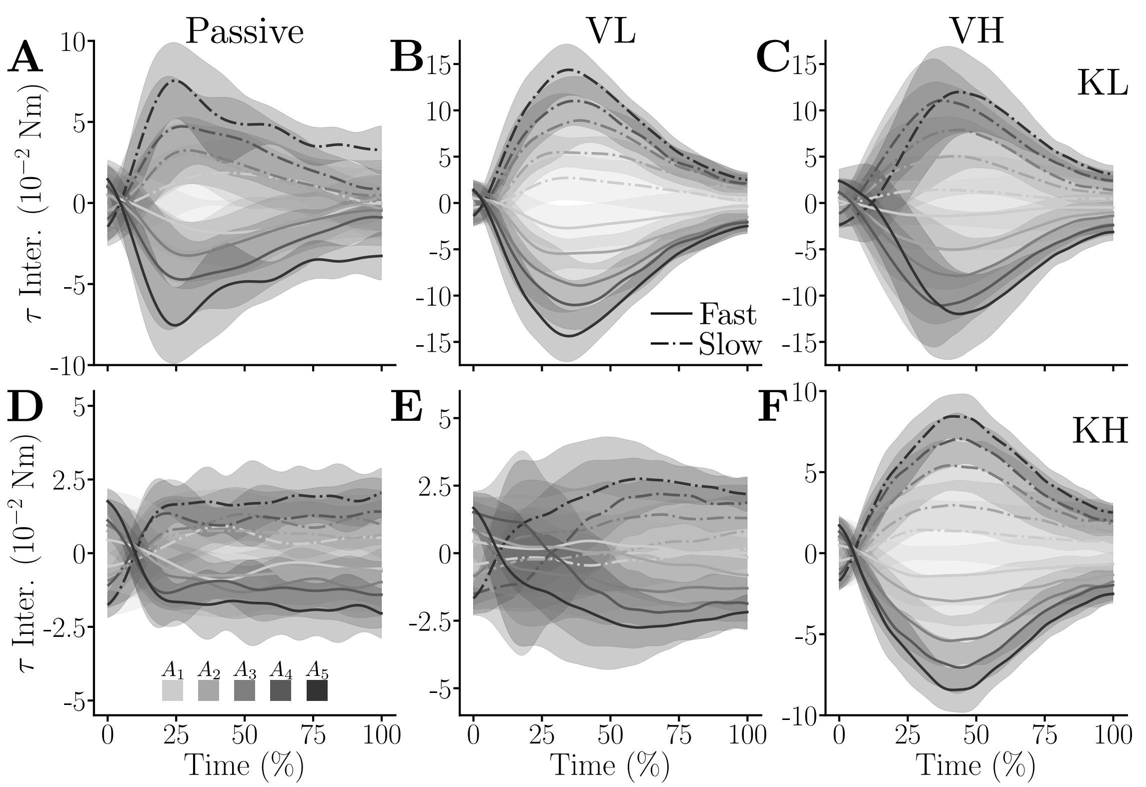


**Figure S.11: Average interaction efforts throughout conditions and targets for dyad D4.** Efforts received by the the more vigorous participant are represented with solid lines and those received by the less vigorous participant are represented with dashed-dotted lines. **A.** KL. **B.** KLVL. **C.** KLVH. **D.** KH. **E.** KHVL. **F.** KHVH.

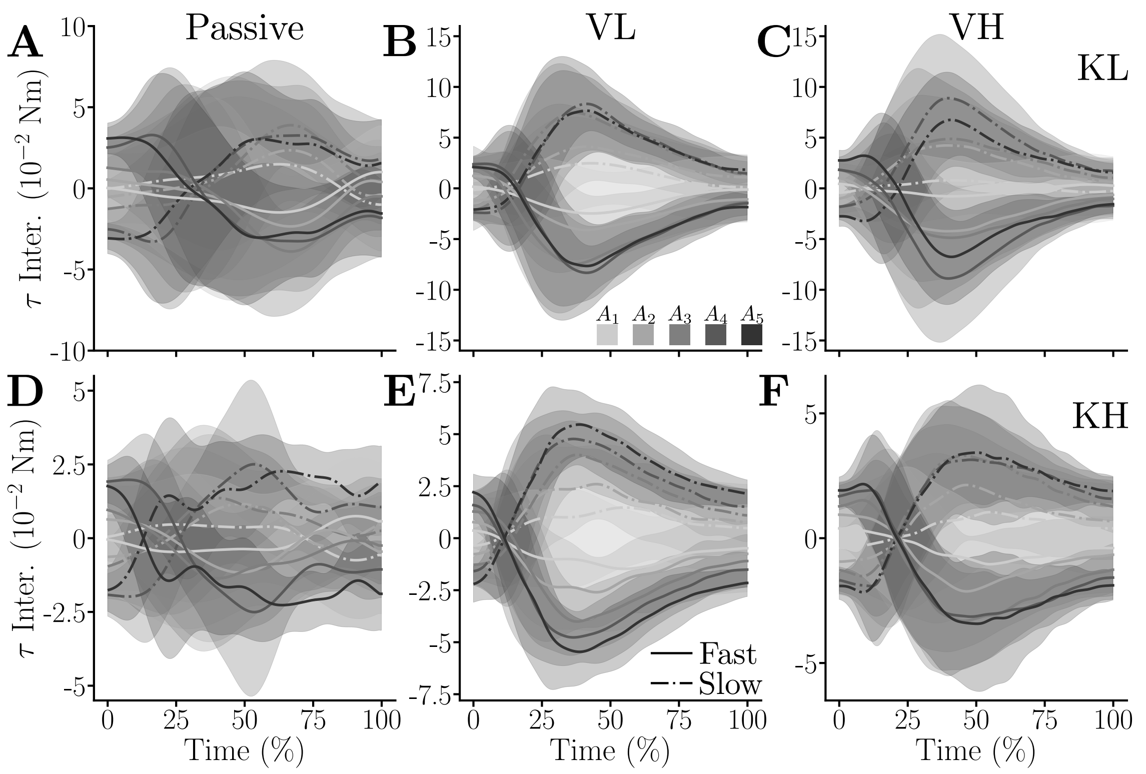


**Figure S.12: Average interaction efforts throughout conditions and targets for dyad D5.** Efforts received by the the more vigorous participant are represented with solid lines and those received by the less vigorous participant are represented with dashed-dotted lines. **A.** KL. **B.** KLVL. **C.** KLVH. **D.** KH. **E.** KHVL. **F.** KHVH.

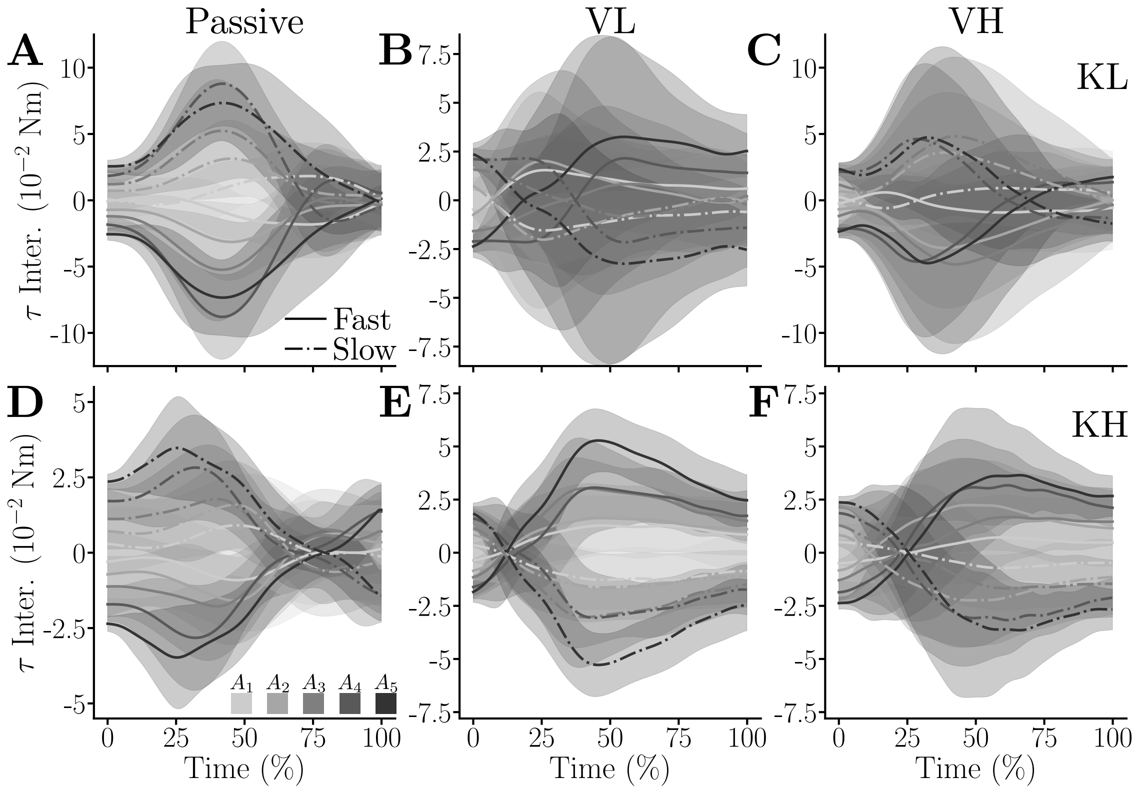


**Figure S.13: Average interaction efforts throughout conditions and targets for dyad D6.** Efforts received by the the more vigorous participant are represented with solid lines and those received by the less vigorous participant are represented with dashed-dotted lines. **A.** KL. **B.** KLVL. **C.** KLVH. **D.** KH. **E.** KHVL. **F.** KHVH.

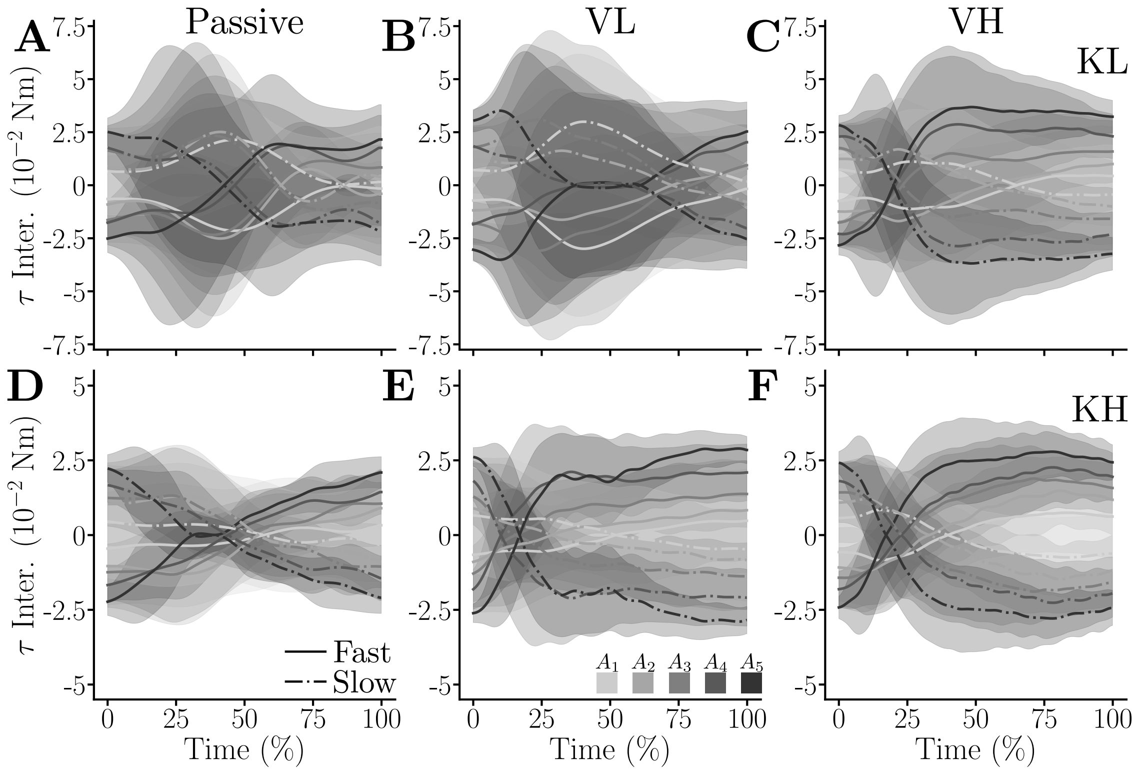


**Figure S.14: Average interaction efforts throughout conditions and targets for dyad D7.** Efforts received by the the more vigorous participant are represented with solid lines and those received by the less vigorous participant are represented with dashed-dotted lines. **A.** KL. **B.** KLVL. **C.** KLVH. **D.** KH. **E.** KHVL. **F.** KHVH.

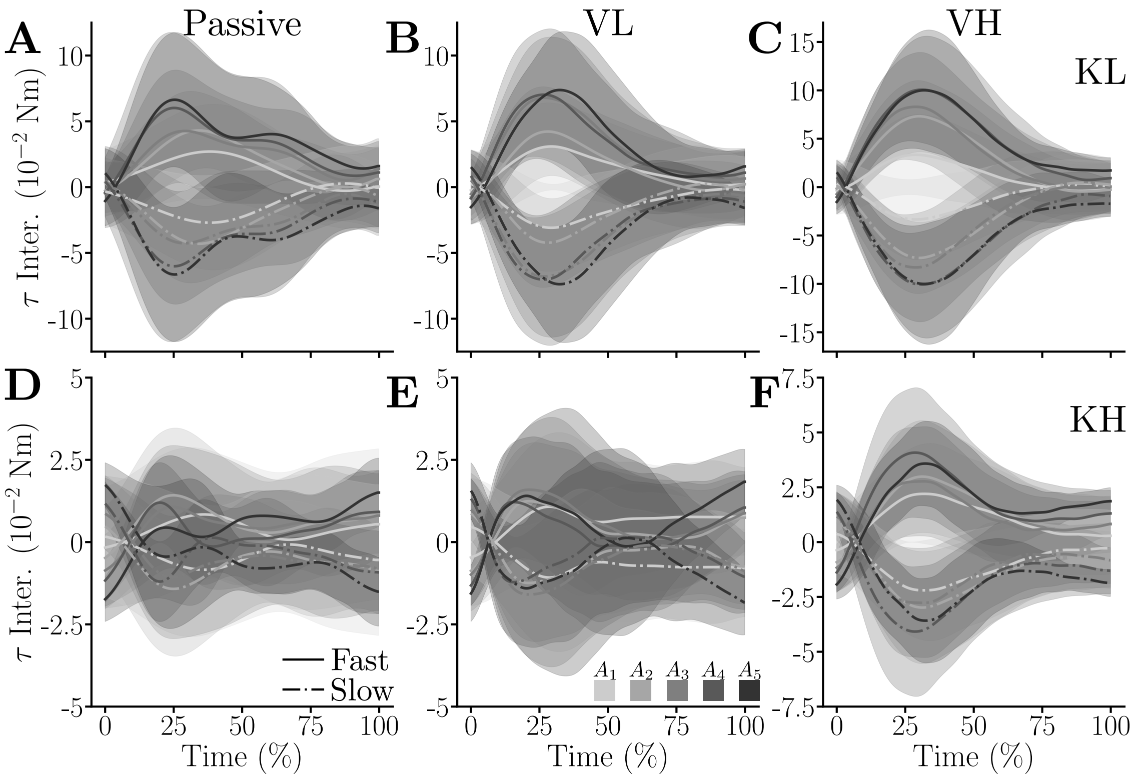


**Figure S.15: Average interaction efforts throughout conditions and targets for dyad D8.** Efforts received by the the more vigorous participant are represented with solid lines and those received by the less vigorous participant are represented with dashed-dotted lines. **A.** KL. **B.** KLVL. **C.** KLVH. **D.** KH. **E.** KHVL. **F.** KHVH.

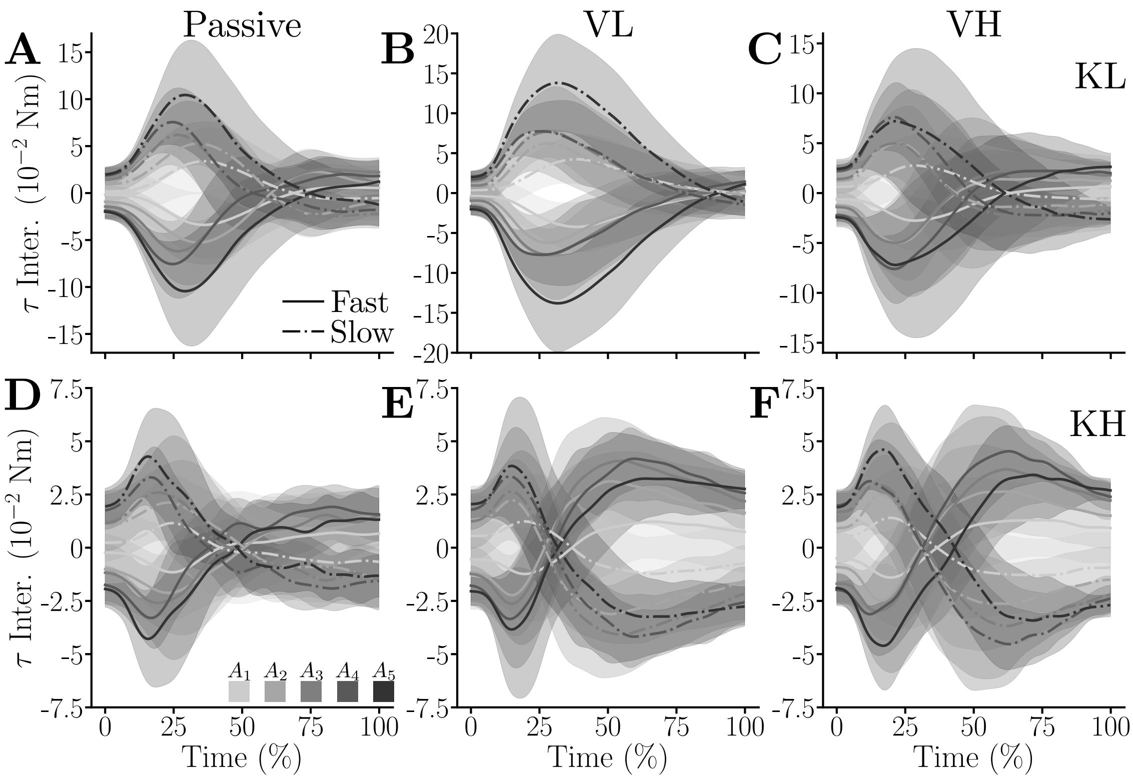


**Figure S.16: Average interaction efforts throughout conditions and targets for dyad D9.** Efforts received by the the more vigorous participant are represented with solid lines and those received by the less vigorous participant are represented with dashed-dotted lines. **A.** KL. **B.** KLVL. **C.** KLVH. **D.** KH. **E.** KHVL. **F.** KHVH.

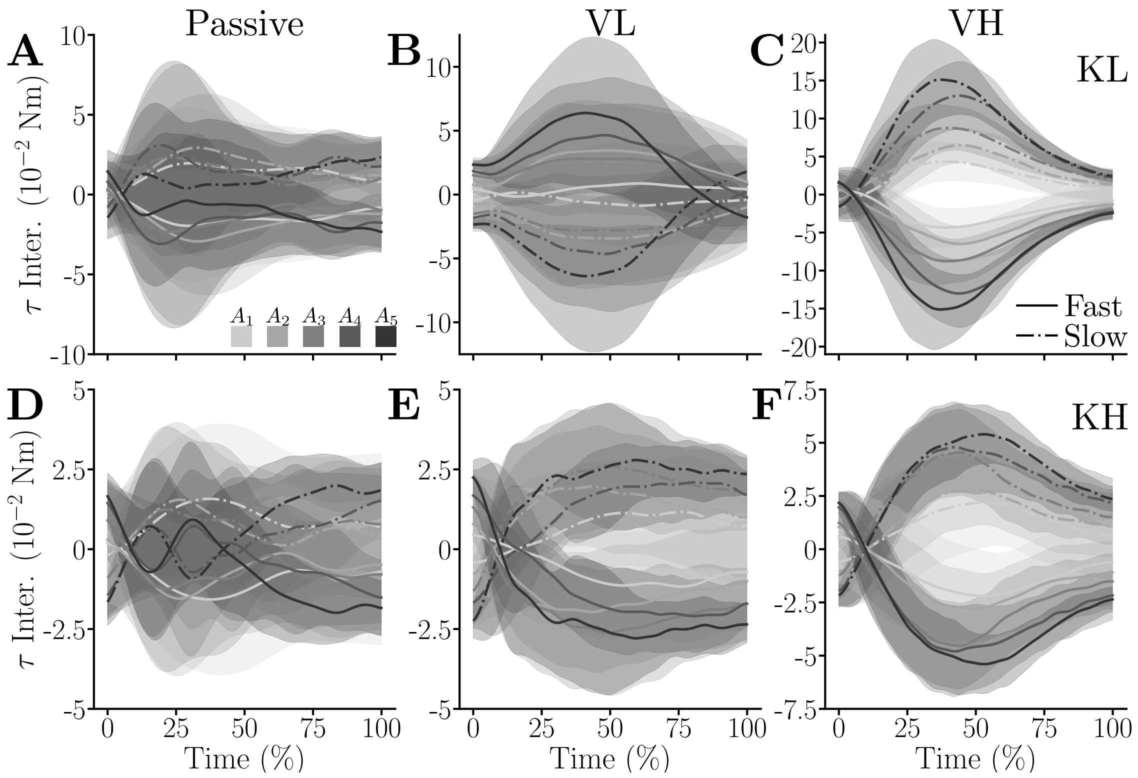


**Figure S.17: Average interaction efforts throughout conditions and targets for dyad D10.** Efforts received by the the more vigorous participant are represented with solid lines and those received by the less vigorous participant are represented with dashed-dotted lines. **A.** KL. **B.** KLVL. **C.** KLVH. **D.** KH. **E.** KHVL. **F.** KHVH.

#### 49 2.4 Complementary results on the human-human modeling

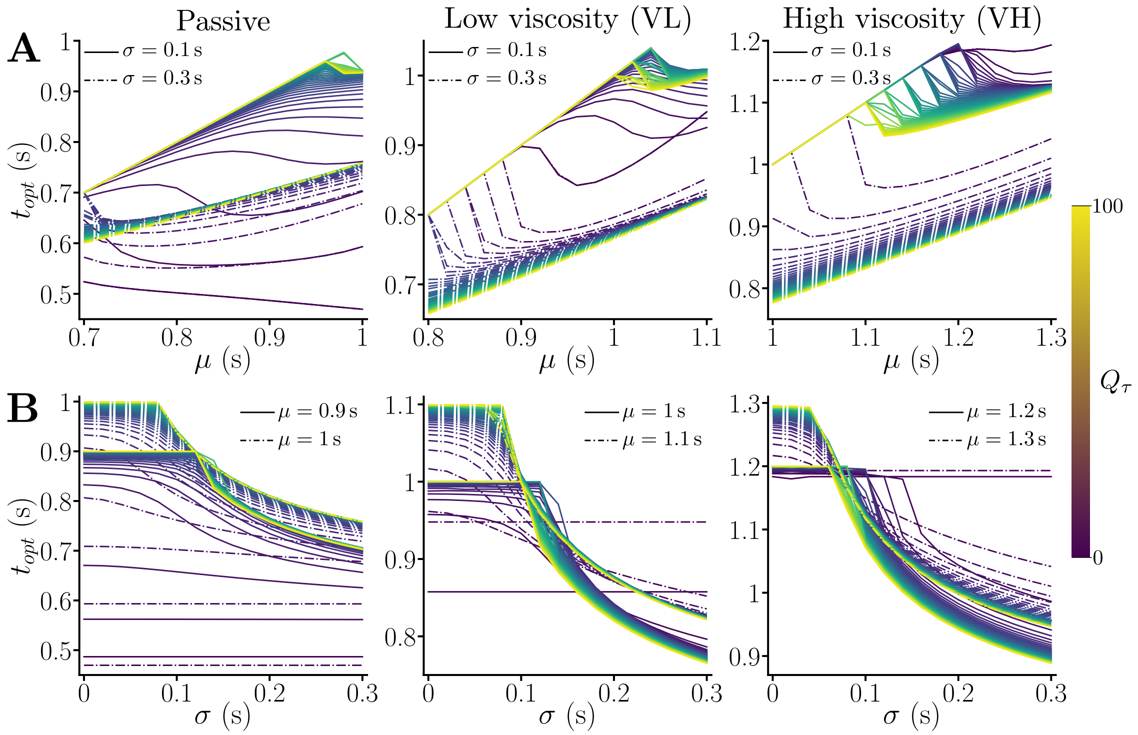


**Figure S.18: Evolution of optimal time predictions for the KL conditions.** The three treated conditions are KL, KLVL, and KLVH. The discretizations of variables spaces was 0.02 s for $\mu$, 0.02 s for $\sigma$, and 2 for $Q_\tau$. **A.** Predicted optimal movement duration for two values of $\sigma \in \{0.1, 0.3\}$ s, and for ranges of average partner movement duration adapted to the condition. **B.** Predicted optimal movement duration for two values of $\mu$ depending on the condition, and for $\sigma \in [0, 0.3]$ s.

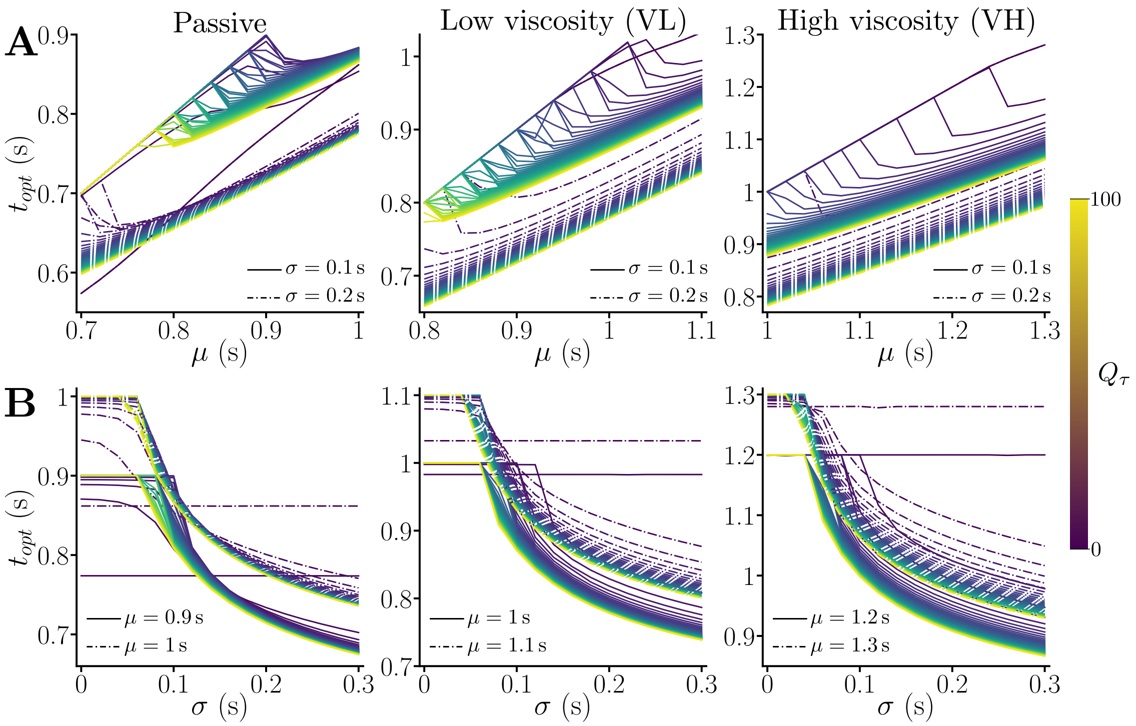


**Figure S.19: Evolution of optimal time predictions for the KH conditions.** The three treated conditions are KH, KHVL, and KHVH. The discretizations of variables spaces was 0.02 s for $\mu$, 0.02 s for $\sigma$, and 2 for $Q_\tau$. **A.** Predicted optimal movement duration for two values of $\sigma \in \{0.1, 0.2\}$ s, and for ranges of average partner movement duration adapted to the condition. **B.** Predicted optimal movement duration for two values of $\mu$ depending on the condition, and for $\sigma \in [0, 0.3]$ s.

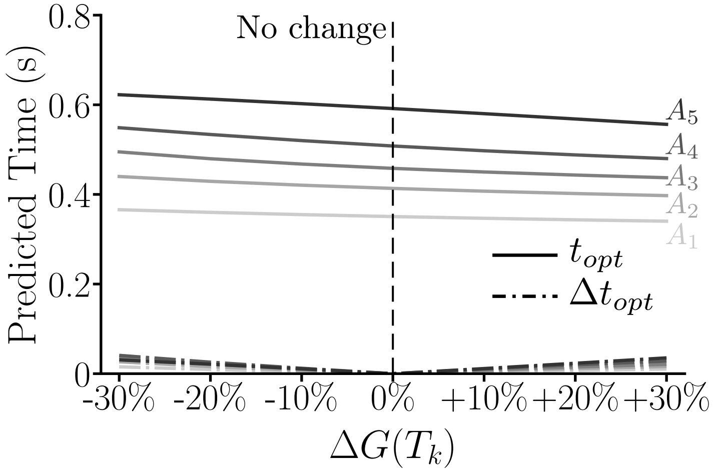


**Figure S.20: Effect of varying the cost of time of the fast partner on predicted movement time.** Solid lines represent predicted times and dashed-dotted lines the absolute change of predicted time when varying $G(T_k)$ with a change in $[-30\%, +30\%]$. These results were obtained for $\kappa = 0.5$ Nm s/rad and the same weighting and uncertainty as in the main paper.

#### 3 After-effects and summary of behavior evolution through the experiment

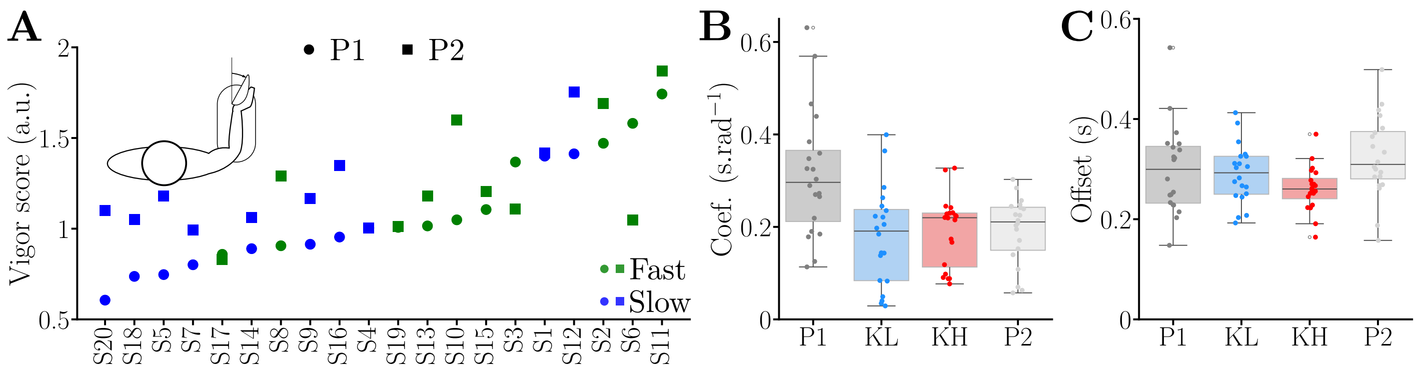


**Figure S.21: Effect of the connection on vigor and the affine amplitude-duration relationship. A.** Distribution of vigor scores in P1 and P2. **B.** Affine coefficient. Main effect of the condition ($W = 0.34$, $p < 10^{-3}$), further confirmed by pairwise comparisons showing that coefficients in P1 are significantly higher than in the other conditions (in the three cases: $p < 2{\cdot}10^{-3}$, Cohen's $D > 1.09$). **C.** Offset. Main effect of the condition ($W = 0.15$, $p = 0.03$), pairwise comparisons showed that offsets were lower in KH than in KL and P2 (in both cases: $p < 0.03$, Cohen's $D > 0.55$).

| | | **Solo** | | | **Dyads** | | | | | | **AE** |
|---|---|---|---|---|---|---|---|---|---|---|---|
| Dyad Id. | Subject Id. | P1 | VL | VH | *High stiffness* KH | VL | VH | *Low stiffness* KL | VL | VH | P2 |
| D1 | S1 | 1.4 | 1.34 | 1.19 | 1.44 | 0.86 | 0.68 | 1.32 | 0.93 | 0.72 | 1.42 |
| | S2 | 1.47 | 1.12 | 1.04 | | | | | | | 1.6 |
| D2 | S3 | 1.37 | 1.15 | 1.17 | 1.21 | 0.82 | 0.64 | 1.17 | 0.79 | 0.64 | 1.11 |
| | S4 | 1 | 0.78 | 0.64 | | | | | | | 1 |
| D3 | S5 | 0.75 | 0.66 | 0.42 | 0.84 | 0.69 | 0.59 | 0.97 | 0.75 | 0.54 | 1.18 |
| | S6 | 1.58 | 1.19 | 0.94 | | | | | | | 1.05 |
| D4 | S7 | 0.8 | 0.67 | 0.61 | 0.94 | 0.64 | 0.56 | 0.87 | 0.64 | 0.57 | 0.99 |
| | S8 | 0.91 | 0.84 | 0.8 | | | | | | | 1.29 |
| D5 | S9 | 0.91 | 0.82 | 0.81 | 1.1 | 0.7 | 0.55 | 1.06 | 0.75 | 0.61 | 1.17 |
| | S10 | 1.05 | 1.16 | 0.54 | | | | | | | 1.6 |
| D6 | S11 | 1.74 | 1.28 | 1.02 | 1.32 | 0.85 | 0.72 | 1.38 | 0.92 | 0.76 | 1.87 |
| | S12 | 1.41 | 1.11 | 0.92 | | | | | | | 1.75 |
| D7 | S13 | 1.02 | 0.7 | 0.56 | 0.94 | 0.59 | 0.43 | 1.06 | 0.64 | 0.47 | 1.18 |
| | S14 | 0.89 | 0.64 | 0.52 | | | | | | | 1.06 |
| D8 | S15 | 1.11 | 1.01 | 0.82 | 1.03 | 0.61 | 0.57 | 1.1 | 0.7 | 0.57 | 1.21 |
| | S16 | 0.95 | 0.98 | 0.75 | | | | | | | 1.35 |
| D9 | S17 | 0.86 | 0.83 | 0.79 | 0.79 | 0.66 | 0.61 | 0.83 | 0.67 | 0.6 | 0.83 |
| | S18 | 0.74 | 0.8 | 0.93 | | | | | | | 1.05 |
| D10 | S19 | 1.01 | 0.73 | 0.65 | 0.8 | 0.56 | 0.49 | 0.68 | 0.53 | 0.59 | 1.01 |
| | S20 | 0.61 | 0.71 | 0.56 | | | | | | | 1.1 |

**Table S.1: Summary of the vigor scores of participants through the experiment.** The vigor scores in the VH and VL conditions (with and without connection) were computed using the average movement durations collected in the corresponding condition without viscous resistance to better show the effects of changes in the dynamics. Vigor scores in the P2 condition were computed using average durations from the P1 conditions to highlight changes after exposure to the human-human connection.
